## Supplementary figures and images for "Isoginkgetin and Madrasin are poor splicing inhibitors"

### Supplementary Figures

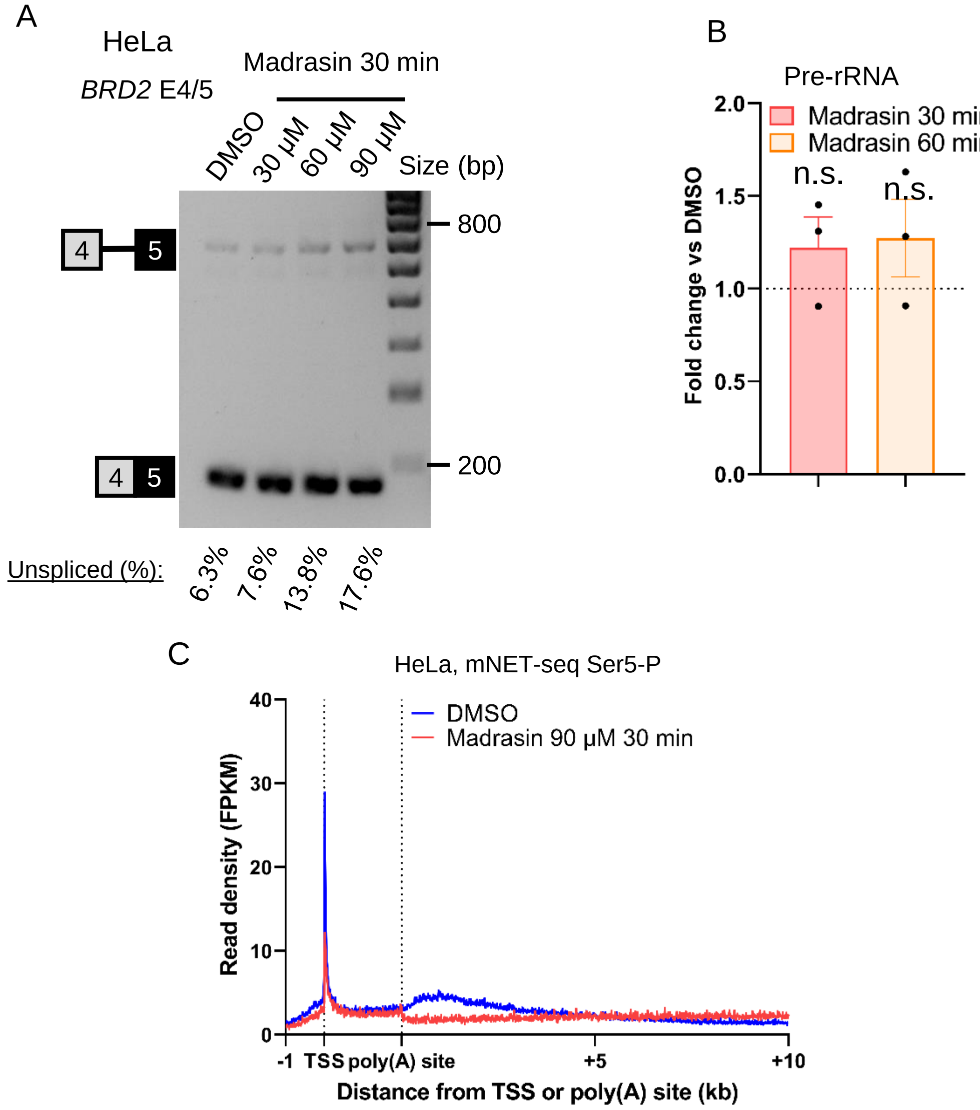

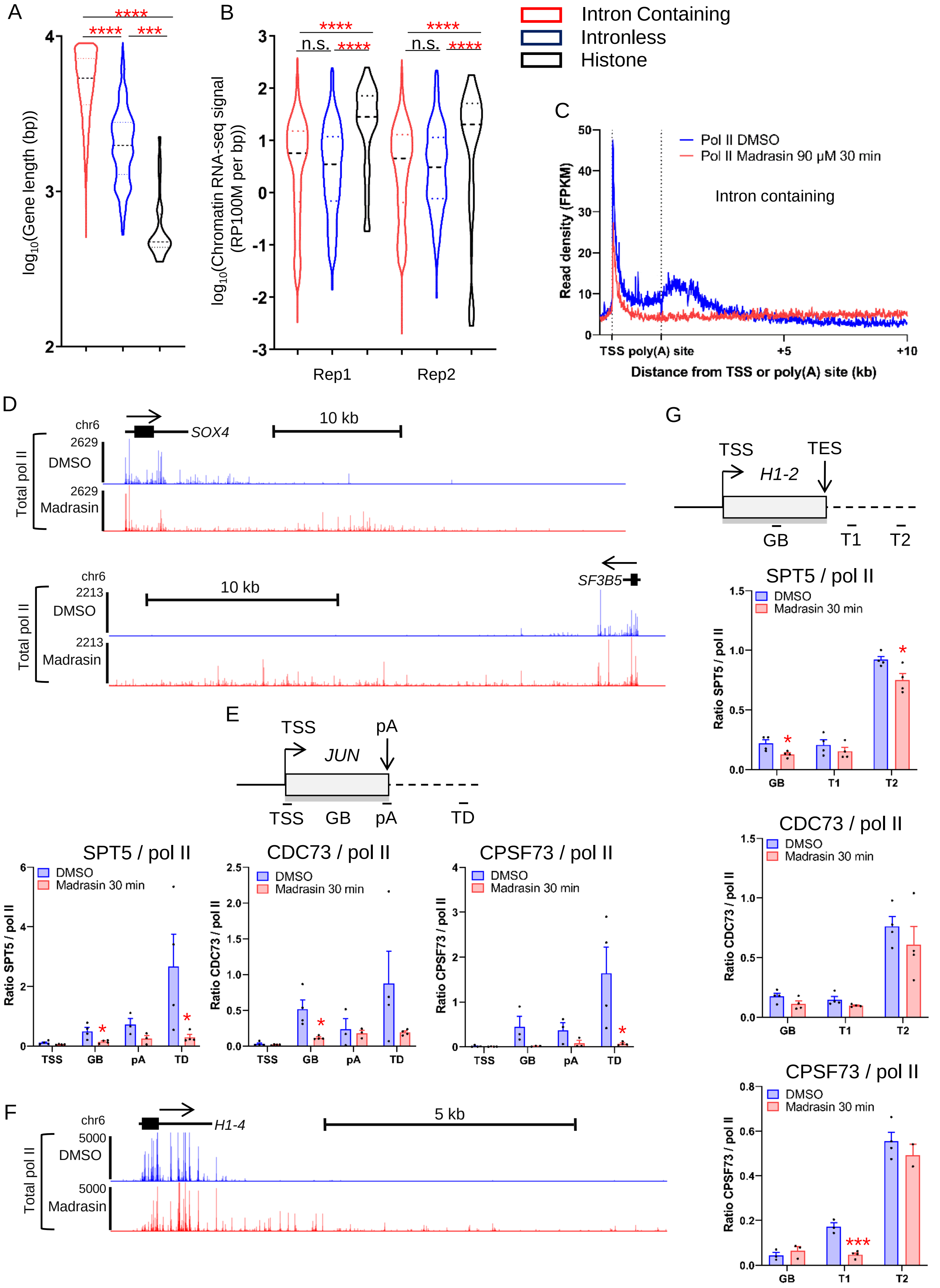

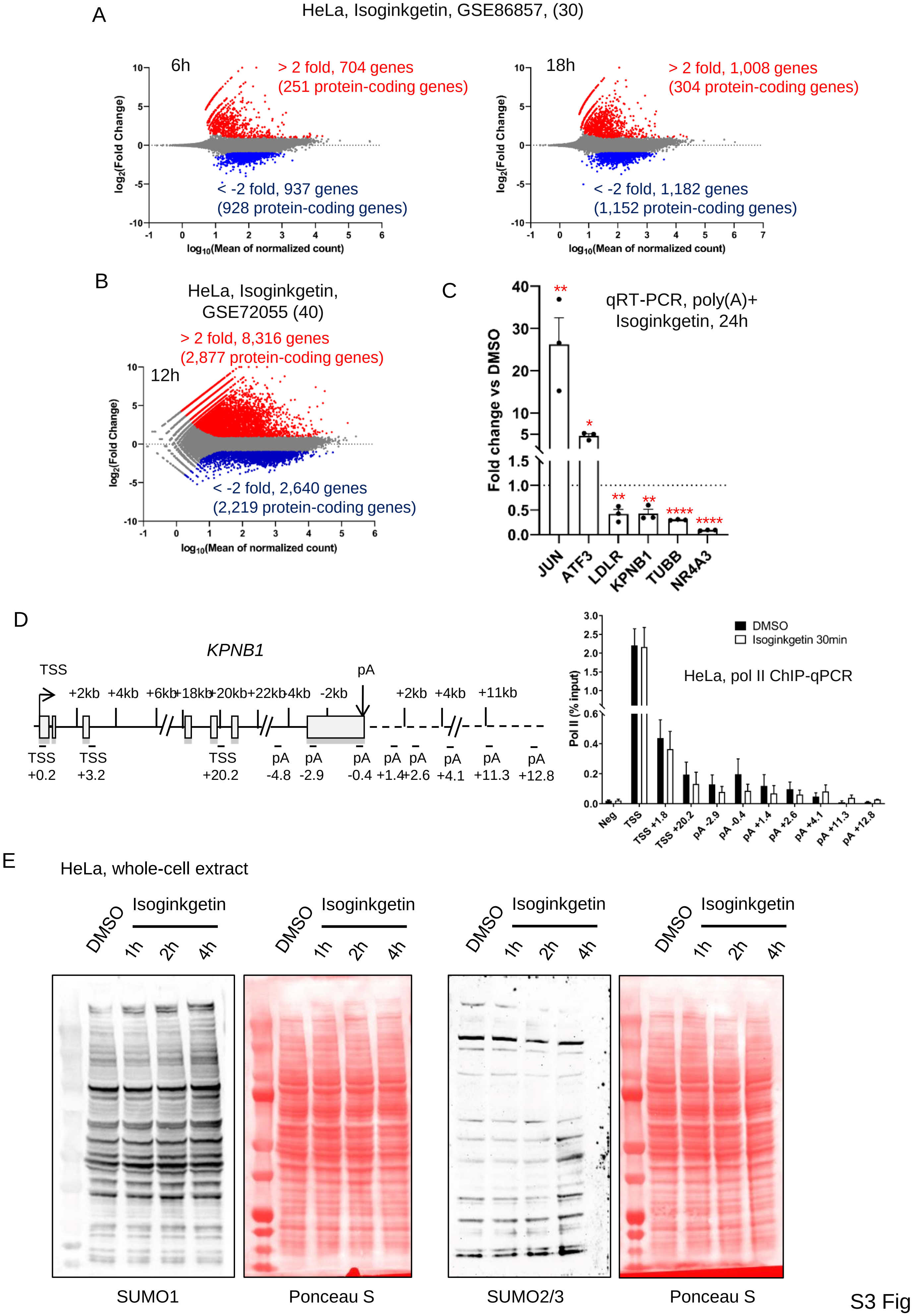

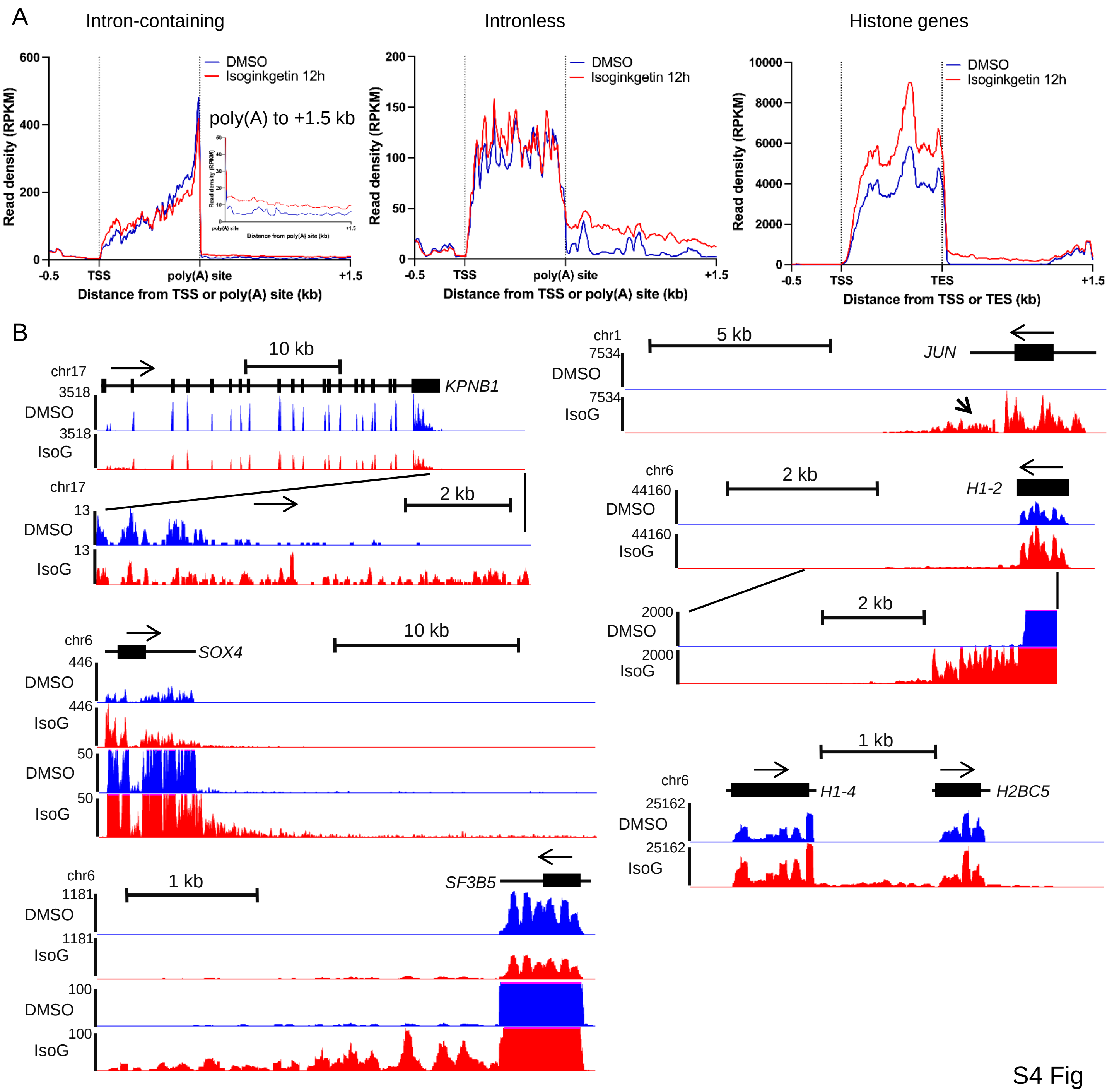
